# Resonant fluctuations of selection pressure exponentially accelerate fitness valley crossing

**DOI:** 10.1101/2022.01.03.474754

**Authors:** Mario E. Di Salvo, Kimberly A. Reynolds, Milo M. Lin

## Abstract

Two functional protein sequences can sometimes be separated by a fitness valley - a series of low or non-functional intermediate mutations that must be traversed to reach a more optimal or refined function. Time-varying selection pressure modulates evolutionary sampling of such valleys. Yet, how the amplitude and frequency of fluctuating selection influence the rate of protein evolution is poorly understood. Here, we derive a simple equation for the time-dependent probability of crossing a fitness valley as a function of evolutionary parameters: valley width, protein size, mutation rate, and selection pressure. The equation predicts that, under low selection pressure, the valley crossing rate is magnified by a factor that depends exponentially on valley width. However, after a characteristic time set by the evolutionary parameters, the rate rapidly decays. Thus, there is an optimal frequency of selection-pressure fluctuations that maximizes the rate of protein optimization. This result is reminiscent of the resonance frequency in mechanical systems. The equation unites empirical and theoretical results that were previously disconnected, and is consistent with time-dependent in vitro and clinical data. More generally, these results suggest that seasonal and climate oscillations could synchronously drive protein evolution at the resonant frequency across a range of organism hosts and timescales. This theory could also be applied to optimize de novo protein evolution in laboratory directed evolution using time-varying protocols.

## I. Main

The mechanisms of natural selection are well established [1–3], yet predicting quantitative properties of evolutionary dynamics remains challenging. The evolutionary dynamics of proteins can be considered using Sewell Wright’s fitness landscape framework [4]. This construct maps sequences (genotype) to fitness (phenotype) following the general rules that more closely related sequences are proximal on the fitness map, and higher fitness sequences occupy peaks on the landscape [5]. Evolutionary trajectories tend to follow neutral or higher-fitness gradients in the mutational fitness landscape, but such trajectories can become increasingly rare over time [6]. As first suggested by Wright [4] and later corroborated by multiple sequence alignment[7], the population may become trapped in a local fitness maximum in which multiple deleterious mutations (constituting a fitness valley) are required to achieve higher fitness. Laboratory directed-evolution experiments also indicate that protein fitness landscapes are “rough,” containing many local fitness maxima separated by fitness valleys [8–11]. Detailed characterization of such a fitness valley was demonstrated for a protein conferring chloroquine resistance to malarial parasites [12], as well as in large-scale directed evolution experiments of a *λ* phage protein [13]. Fitness valleys can explain why directed evolution of *de novo* enzymes typically plateaus at much lower performance levels than that of natural enzymes [14–16]. Crossing fitness valleys is therefore a general rate-limiting process for protein design and evolution.

It has long been appreciated that valley crossing can be catalyzed by mechanisms that decrease the fitness cost of intermediates. For example, following gene duplication, one of the gene copies may tolerate intermediate mutations while the other copy is maintained by purifying selection [17]. External (environmental) fluctuations can also serve to temporarily relax selection pressure. Dramatic fluctuations in selection pressure may occur in response to incremental changes in the physical environment (such as nutrient availability or climate) [18–21]. The temporal frequency of fitness fluctuations spans many orders of magnitude depending on the timescales of, e.g., ecological versus climate change, and are consistent with evidence of the temporal heterogeneity of extinction and speciation [22, 23]. To connect environmental change to valley crossing, work has been done in the case in which the fitness function happens to change such that deleterious mutants become momentarily more fit (i.e. moving the peaks in the fitness landscape) [24–26]. These results are contingent on the specific shape of the changing fitness landscape, which becomes difficult to enumerate and mathematically generalize due to the large number of dimensions involved. In contrast, the regime in which the fitness landscape is dynamically rescaled by a single factor due to the time-varying importance of the protein to organism fitness has not been quantitatively explored.

Most existing work considers valley crossing of the entire genome (i.e. the fate of species) in a static fitness landscape. In this case, models typically focus on one or a few dominant trajectories within the high-dimensional sequence space, with the assumption that other possible mutations are either decoupled from the evolutionary process of interest or are suppressed by strong purifying selection pressure on the other genes in the genome. In this case, an asexual population can cross fitness valleys by either sequential fixation of the lower fitness intermediates or by simultaneous fixation (stochastic tunneling), depending on population size [27–29]. The valley-crossing probability has been derived in both the large population limit [30], as well as for intermediate and small asexual [31] and sexual [32] populations in which valley-crossing is dominated by the discrete fluctuations underlying genetic drift. This work differs from previous results in two ways. First, we treat the individual protein sequence as the evolutionary unit rather than the genome. Because different genes are under different kinds of selection pressure due to multiplicity of gene functions and regulatory relationships, a change in the environment will change the genome fitness landscape in a non-uniform (and even non-monotonic) manner. In contrast, because proteins generally perform a set of fixed tasks, the change in the environment will scale the protein fitness landscape in an approximately uniform manner proportional to the importance of the protein’s function in the new environment relative to the original environment. This property simplifies the problem by reducing an otherwise case-specific high-dimensional environmental variable into a scalar selection amplitude. This also means that this theory is concerned with optimization of a fixed task, rather than the process of protein differentiation into new tasks (although this theory is applicable to protein differentiation under appropriate conditions; see below). Second, because we are interested in the weak selection regime facilitating mutational diffusion, we do not assume that all mutations in “off-target” sites (amino acid positions) are suppressed by purifying selection. Instead, we explicitly consider the (reversible) evolutionary dynamics during weak selection of all protein sites that would be sensitive to mutation under strong selection. Together, these two features constitute a modeling paradigm of evolutionary dynamics in a high-dimensional sequence space modulated by a single time-varying selection amplitude.

We derive a simple mathematical expression for the valley-crossing probability of protein evolution under varying modulation of selection frequency and amplitude. We show that the rate of valley crossing can vary by many orders of magnitude depending on the timescale over which selection pressure is modulated as well as the amplitude of the modulation. Specifically, we calculate the probability that *w* sites that have reciprocal sign epistasis acquire a specific set of mutations that lead to a new fitness peak during a period in which the selection pressure on the number of function-sensitive sites, *L*, are relaxed (See Fig. 1). Intuitively, the optimal time should not be too short so as to allow for enough time to acquire the *w* mutations, but should not be too long such that the *L* sites explore too much of the exponentially large (e.g. 20^*L*^) mutation space, thereby rendering the target mutant improbable. The optimal tradeoff between these two mechanisms occurs at the “resonant” period *τ*_res_, which depends on the constituent parameters. For time-varying selection pressure, if selection is turned on and off at the resonant frequency 1/*τ*_res_, then the rate of traversing a fitness valley is amplified exponentially as a function of the fitness valley size.

**Figure 1:**
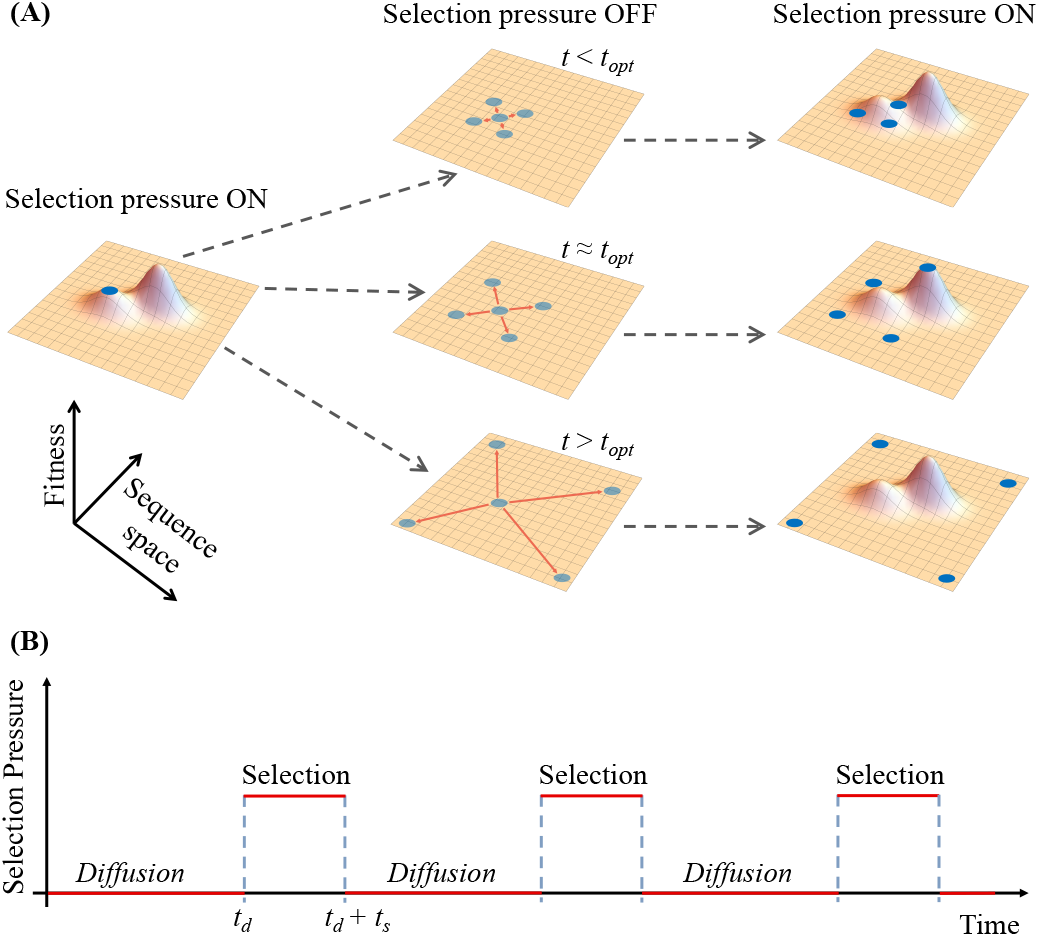
Escaping local fitness maxima via environmental fluctuations. **(a)** Schematic representation of a fitness landscape: When selection pressure is relaxed, the population that is trapped diffuses in sequence space. The probability of crossing a fitness valley is maximized for an optimal relaxation time window. **(b)** Diagram of an idealized selection pressure fluctuation. Each fluctuation period comprises a diffusion time *t*_*d*_ during of weak or no selection and a selection time *t*_*s*_ during of strong selection.

## II. Results

### Protein fitness valley-crossing equation under weak selection

We will define mutation and selection on the protein sequence, where *μ* denotes the effective mutation rate from one amino acid to another specific amino acid per copy of the gene encoding the protein. Using this definition of *μ*, the alphabet size *n* = 20. It may sometimes be convenient to consider mutations between specific gene sequences, in which case *n* = 4. Note that the values of *μ* and *N* will also depend on alphabet choice. In the analytic model, we treat *μ* as a single parameter reflecting the mean mutation rate between pairs of amino acids. In our simulations, we use sequence-dependent empirical values of *μ*. We model the “host” of the protein sequence to be an asexual organism, and define the population size *N* to be the copy number of the gene encoding the protein. This is consistent with the predominant environment of proteins on evolutionary timescales (e.g. prokaryotes), as well as in hosts with fast enough mutation rate *μ* for which protein valley crossing is relevant on physiological timescales (e.g. viruses, cancer cells, and cell lineages designed for accelerated laboratory evolution [33]). Therefore, we adopt a master equation approach that is meaningful if *μN* is not much smaller than 1. In addition, this theory focuses on the valley crossing to reach a specific target sequence with high fitness. In reality, the probability of reaching any new peak must take into consideration the multiplicity of valley-separated peaks.

Consider the case in which the protein is at a local fitness maximum. If the function of the protein is under strong selection, define *L* to be the number of amino acid positions within the protein that are sensitive to mutation. In this case, mutation of any of the *L* sites would be deleterious. Note that, depending on the protein fold and function, *L* could be the entire sequence, or a subset of amino acids composed of, e.g., the active site, the protein core [34], and/or co-evolved allosteric sectors [35]. Assume that there is a fitness valley of size *w* − 1. *w* corresponds to a subset of the *L* sites, such that mutation of all *w* sites to specific amino acids would lead to a higher fitness mutant, but mutations of any other subset of *L* is deleterious. The population size is assumed to be fixed. During relaxed selection, in each generation, every site (i.e. amino acid position) has probability *μ*(*n* − 1) of mutation as well as probability *α* that, if this site is not the wild-type mutant, it will revert back to the wild-type due to weak selection pressure. Therefore, during weak selection we assume the simplest possible model in which mutation at each of the *L* sites leads to an additive fitness cost *α* relative to wild type. Competition between *μ* and *α* maintains standing variation. The probability per individual, *P*_*w*_(*t*), of achieving the target mutations at *w* sites while not mutating the remaining *L* − *w* sites in time *t* is: *P*_*w*_(*t*) = *p*(*t*)^*w*^[1 − (*n* − 1)*p*(*t*)]^*L*−*w*^, where *p*(*t*) is the probability that a site will have mutated to a specific amino acid in time *t* during relaxed selection. The key quantity is therefore *p*(*t*), which is a result of mutation and reversion. Although exact values of mutation and reversion rates are sequence dependent (See simulation results to follow), we first obtained the analytic solution of *p*(*t*) assuming constant characteristic values of *μ* and *α* (see details in the supplementary materials):

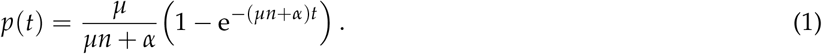

Therefore, the mutation-reversion equation for crossing a valley of size *w* − 1 in time *t* is:

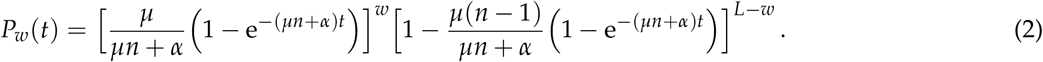

Note that *P*_*w*_(*t*) must be multiplied by the population size to give the probability of achieving *w*-level innovation. We also assume that achieving all *w* mutations leads to a significant fitness advantage for the protein host when strong selection is turned on. Therefore, if the valley is crossed, there is a reasonable chance that the mutant will fix in the population. Note that Eq. 2 is also applicable to the case of crossing a fitness valley in response to a new fitness landscape due to changing functional requirements (e.g. enzymatic activity for a slightly different metabolite source or toxin) as long as the population starts in a local fitness optimum of the original landscape, and fluctuating selection also occurs in the original landscape prior to introduction of the new landscape conditions. This scenario is relevant for the evolution of new functional targets, such as the case for bacterial resistance to new antibiotics (See below).

Eq. 2 explicitly relates the probability of crossing a valley of width *w* − 1 in time *t* as a function of the elementary parameters of mutation rates, reversion rates, and mutational sensitivity. Starting from this expression for *P*_*w*_(*t*), we show in the following sections that simple relations can be derived to address multiple evolutionary questions under static and dynamic environmental conditions.

### Limits on mutation rate, time, and population size

By optimizing Eq. 2 with respect to different constituent parameters, we can derive limits on maximum mutation rate, characteristic waiting times, and minimum population size containing a valley-crossing mutant. Where available, we show that these limits are consistent with experimental data and/or theoretical results that were previously disconnected. These limits show how Eq. 2 can provide unifying intuition about the constraints of evolutionary optimization.

For the maximum mutation rate, consider the case *t* = 1, *w* = 1 and *α* = 0; *P*(1) in this case corresponds to the probability of mutating to the nearest neighbor in sequence space during one generation. If we define *L* to correspond to the genome length, then the value of *μ* that maximizes *P*(1), denoted *μ*_*max*_, corresponds to the mutation rate that maximizes the acquisition of single mutants under continuous selection. Differentiating Eq. 2 with respect to *μ* under these conditions, we obtain:

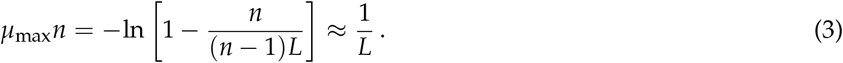

where *μ*_max_*n* is the per-site probability of observing any mutation per generation. If *μ > μ*_*max*_, the number of single mutants decreases because of the prevalence of multiple mutations in one generation: the mutation rate is so high that evolution is no longer a gradient search even under continuous selection. This “error threshold” relation is empirically observed [36, 37]. For example, the genome length of norovirus, one of the fastest mutating viruses, is 7.7 kilobases. The norovirus mutation rate is 1.5 × 10^−4^ per nucleotide per infection, indicating that norovirus mutates at the upper limit allowed by Eq. 3 [38].

The maximum of *P*_*w*_(*t*) occurs at the most likely time that a protein will cross a valley of size *w* − 1, which we denote by the characteristic time *t*_char_. If *α << nμ*, (See Supp. Materials for details):

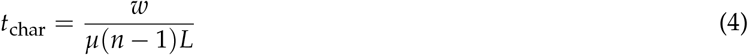

where the denominator *μ*(*n* − 1)*L* is the null mutation rate per locus. Therefore, in the context of weak selection following gene duplication, if the duplicated gene does cross the fitness valley, it should occur on timescale of *t*_char_. This is consistent with Watterson’s prediction that the mean time to nonfunctionalization of the duplicated gene is approximately the reciprocal of the null mutation rate per locus [39, 40], and ranges from a few to tens of millions of years across eukaryotes [41].

In the long-time limit, due to the finite size of the sequence space, *P*_*w*_(*t*) will reach a nonzero asymptotic value, corresponding to a steady-state “persister” population of the valley-crossing mutant present in the standing variation around the wild type. For example, the persister mutant could correspond to an enzyme variant that would enable its survival under conditions of strong selection that is poorly tolerated by the wild type. The persister probability is maximized if the reversion and mutation rates have the ratio: 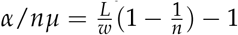. At this optimal ratio of selection to reversion, there is a threshold population size *N*_latent_ such that there will be a persister subpopulation (See Supp. Materials for details):

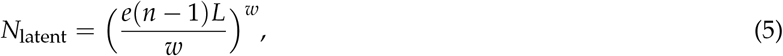

where *e* is Euler’s number. Since *L* typically varies between tens to hundreds, this means that populations of at least hundreds of thousands to tens of millions are necessary to support a persister double mutant at the optimal value of *α*/*nμ*. For *α*/*nμ* that is optimal for triple mutant persisters, a minimum population size of tens of millions to tens of billions (depending on *L*) are necessary to support a given triple mutant persister.

### Universal mutational response curve

Eq. 2 can be used to directly fit data from time-resolved evolutionary processes. We first analyzed data from *in vitro* experiments in which a mutant library of TEM1 beta lactamase was tested against the novel antibiotic cefotaxime [42]. Cefotaxime survival probability was measured following increasing rounds of random mutagenesis followed by selection under different concentrations of ampicillin. Because the starting population was optimized for the original function (ampicillin resistance), and relaxed selection was also under the original function, Eq. 2 is applicable for describing the probability of reaching a target mutant relevant for the new function (cefotaxime resistance). A single mutant was sufficient to confer resistance to cefotaxime (*w* = 1). Therefore, the two fitted parameters are *L* and *μ* (solid lines in Fig. 2A-C). The fitted mutation rates of 1.9 × 10^−3^, 0.8 × 10^−3^, and 0.4 × 10^−3^ under zero, low, and high ampicillin concentration, respectively, agree with the measured effective mutation rates within experimental error [42]. *L* corresponds to the number of sites (nucleotides) that are important for cefotaxime resistance, and was found to range from 112-136 depending on concentration of ampicillin selection in the mutagenesis step; therefore, only about fifteen percent of the TEM1 gene is strongly constrained by cefotaxime resistance. Due to the high mutation rate and multiplicity of ampicillin-tolerant mutants, the best-fit reversion rate *α* is zero. For this special case that *α* = 0, *w* = 1, and mutations per gene are rare within the time window of the experiment, Eq. 2 reduces to a simple “memoryless model” in which the probability of acquiring a deleterious mutation at each site is constant in time and the number of acquired mutations is proportional to time (dashed lines coincide with solid lines in Fig. 2A-C) [42].

**Figure 2:**
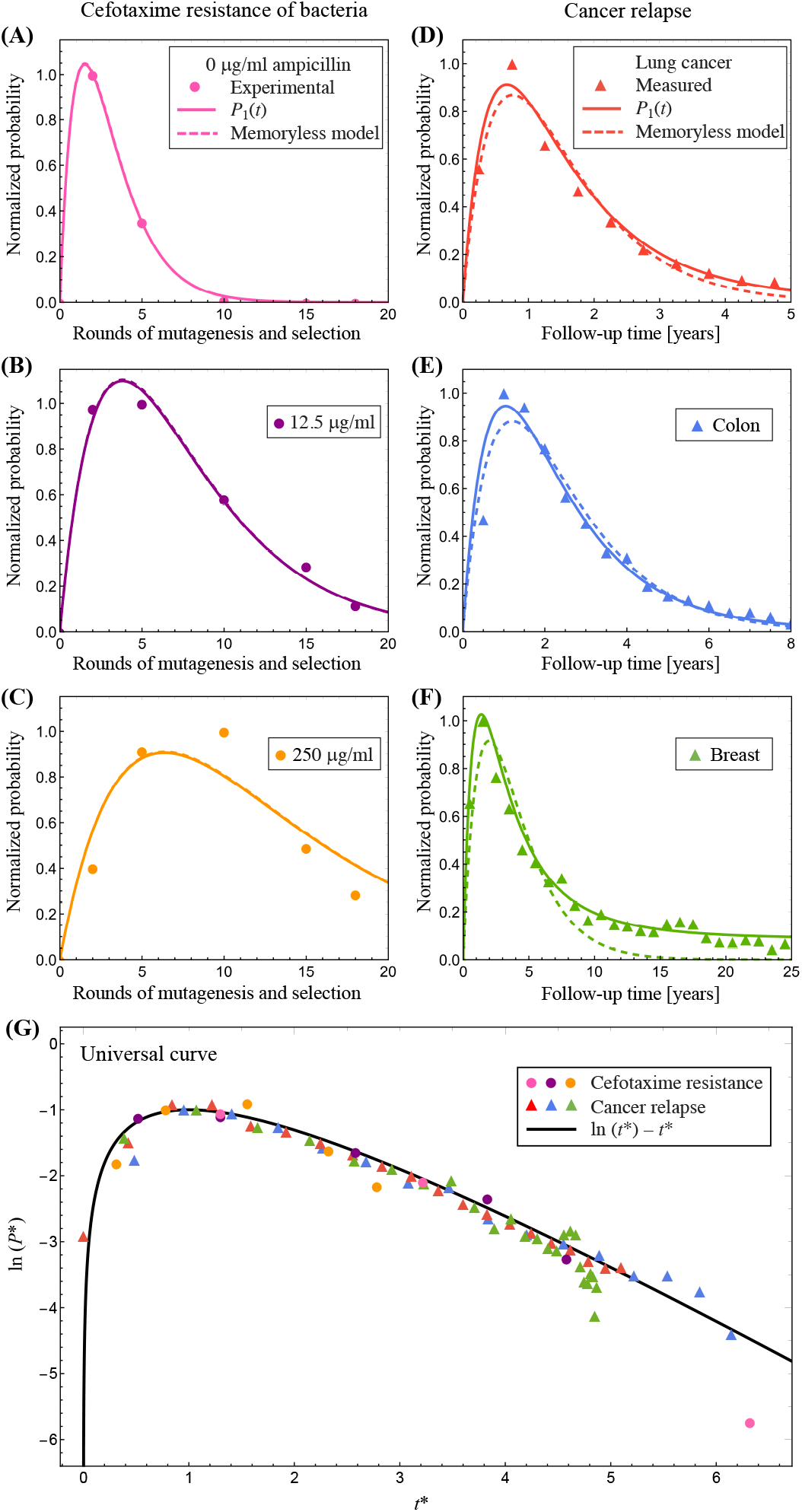
A-C: Cefotaxime resistance of bacteria: circles are experimental data from Ref. [42], continuous lines are a fitting with Eq. 2 (setting *w* = 1) and dashed lines correspond to Eq. 5 in Ref [42]. D-F: Cancer relapse: triangles are measurements from Ref [43–45] continuous line is a fitting with Eq. 2 (setting *w* = 1) and dashed lines correspond to the case *α* = 0. G: Universal function: measurements from the previous references were mapped using the transformation *t*^*^ and *P*^*^ with the values of the parameters obtained by the fitting curves, and the continuous black line is Eq. 6.

Eq. 2 can be used to model evolutionary processes for which the underlying mechanisms are not directly measurable, and for which reversion to the wild-type is non-negligible. A broad class of examples include the evolution of drug resistance in persister sub-populations protected by states of dormancy or residing in microenvironments with limited drug penetrance [46]. We compiled data for cancer relapse time as an example of a mutational response process in which the reversion rate *α* is most likely not negligible compared to the mutation rate. Consider a phenomenological model in which a surviving population of tumor cells are protected from cyclic administration of chemotherapy and can undergo subsequent biased diffusion away from the wild type. In the case of a single resistant mutation (*w* = 1), which is (almost) neutral in the protected tumor but beneficial when the tumor proliferates during cancer relapse, *P*_*w*_(*t*) is the waiting time probability distribution for relapse. We fit three sets of experimental measurements on cancer relapse to Eq. 2 (Fig. 2D-F). The data sets correspond to lung, colon and breast cancer and were modified from the original figures by normalizing the probability to its maximum [43–45]. The fitted mutation rates of 3.9, 2.5, and 2.3 mutations per kilobase per year is in excellent agreement with reported “high tumor mutation burden” rates reported in the literature for lung, colon, and breast cancer, respectively [47–49]. As with the beta-lactamase data, the fits were also performed with the memoryless model. In this case, the memoryless model underperformed compared with Eq. 2. This is mainly due to the importance of the reversion rate *α* in maintaining long-lived standing variation around the wildtype. For breast cancer especially, this explains the signature characteristic of persistent relapse even decades after remission (Fig. 2F). These data suggest that Eq. 2 can be used to model diverse evolutionary processes occurring on different timescales, and that such processes have a characteristic mutational response curve despite variability in the mechanistic details. We can formalize this characteristic shape by defining the transformation 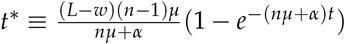, and *P*^*^ ≡ (*n* − 1)(*L* − *w*)*P*_*w*_. In these transformed coordinates, Eq. 2 predicts the universal shape of the mutational response function:

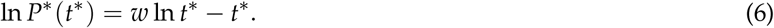

When mapped to Eq. 6, both sets of data collapse to the universal curve (Fig. 2G) with *w* = 1.

### Resonant environmental fluctuations exponentially enhance valley-crossing

Thus far, we analyzed the valley-crossing probability under static conditions of mutation and weak selection. We next explored the implications of Eq. 2 in the case of fluctuating selection pressure that alternates between strong and weak selection. Because convergence to the steady-state distribution under strong selection occurs on much shorter timescales than mutational diffusion under weak selection, we focus on the latter timescale. We define the rate of evolutionary innovation as the valley-crossing probability per generation under weak selection:

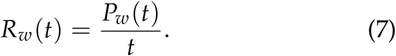

Because we are now interested in the valley-crossing rate (i.e. probability per time), the relevant temporal variable is the period of a selection fluctuation, whereas the total timescale could be a multiple of this period. As with *P*_*w*_(*t*), *R*_*w*_(*t*) should be multiplied by population size to calculate the valley-crossing rate of the population. The maximum of *R*_*w*_(*t*) occurs at the optimal time period of weak selection in order to achieve the maximum rate of crossing an epistatic valley of width *w* − 1 per time; we denote this the “resonant” time *τ*_res_. The optimal valley-crossing protocol would be to alternate periods of strong selection spaced by periods of weak selection of duration *τ*_res_ (diffusion phase in Fig. 1B). In this case, we can characterize the environmental fluctuation frequency as being resonant with the valley-crossing frequency of the protein.

The resonance time will take two different forms depending on whether mutational diffusion dominates over selection back to the wild-type. We call these the mutation-controlled and the reversion-controlled regimes, respectively, and we describe them in turn.

#### mutation-controlled regime

If the mutation rate dominates over reversion to the wildtype, *α << μ*(*n* − 1)*L*, we can ignore *α* to obtain the simplified resonance time (See Supp. Materials for details):

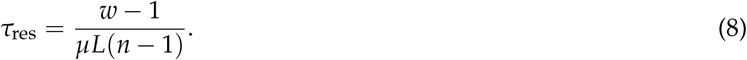

*R*_*w*_(*t*) is plotted as a function of *t* for different epistatic orders *w* in the mutation-controlled regime using E. coli mutation rate (Fig. 3). Note that *τ*_res_ differs from *t*_char_ by the difference of one in the numerator. This is consistent with the expectation that for single mutations (*w* = 1), *τ*_res_ = 0 (i.e. continuous selection pressure corresponds to the fastest innovation rate) because *R*_1_(*t*) is a monotonic function. Note that, in contrast, *P*_1_(*t*) is non-monotonic (See Fig. 2) because there is a characteristic mutation time even if there is no fitness valley. For *w >* 1, *R*_*w*_(*t*) is maximal when *t* = *τ*_res_. Since *τ*_*res*_ ∝ *μ*^−1^, the mutational probability at resonance 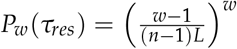 does not depend on the mutation rate of the organism. Therefore, the longest epistatic valley *w* − 1 that can be crossed per resonance period depends only on the population size and *L* (Fig. 4). Substituting Eq. 8 into Eq. 7, the maximum enhancement *ϵ* of the rate of crossing a fitness valley of length *w* − 1 over that of continuous selection is:

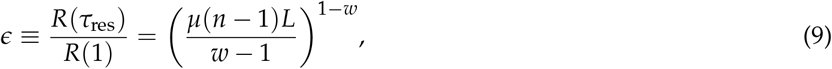

where *R*(*τ*_*res*_) and *R*(1) are the valley-crossing rates under resonant selection pressure and continuous selection pressure, respectively. The resonant rate enhancement is maximized for wide fitness valleys and low mutation rates. The rate enhancement grows exponentially with the length of the epistatic valley, with the base of the exponent being controlled by the mutation rate.

**Figure 3:**
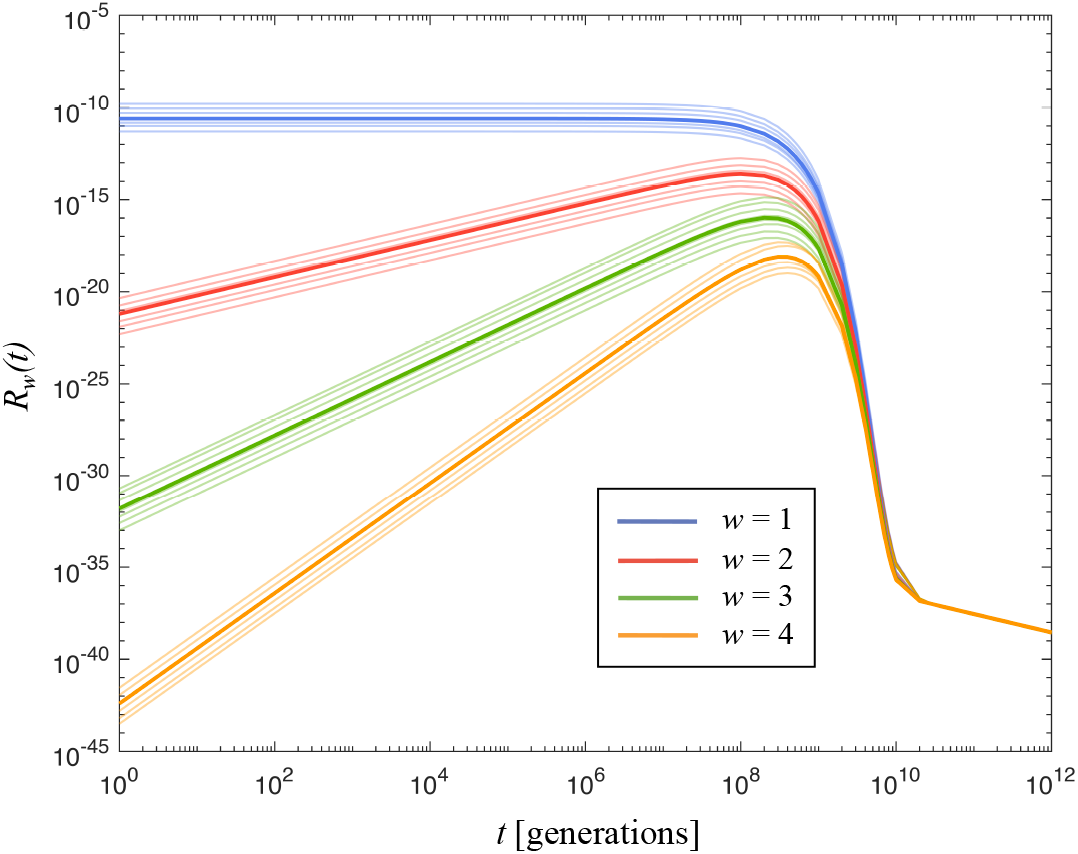
Bacterial protein valley-crossing rate as a function of relaxed selection period. *w* − 1 is the valley size. Thick lines are the analytical results in the mutation-controlled regime using the average E. coli mutation rate between all pairs of amino acids. Thin lines correspond to heterogeneous mutation rates between different pairs of amino acids obtained from the empirical PAM matrix. *L* = 20 in these simulations.

**Figure 4:**
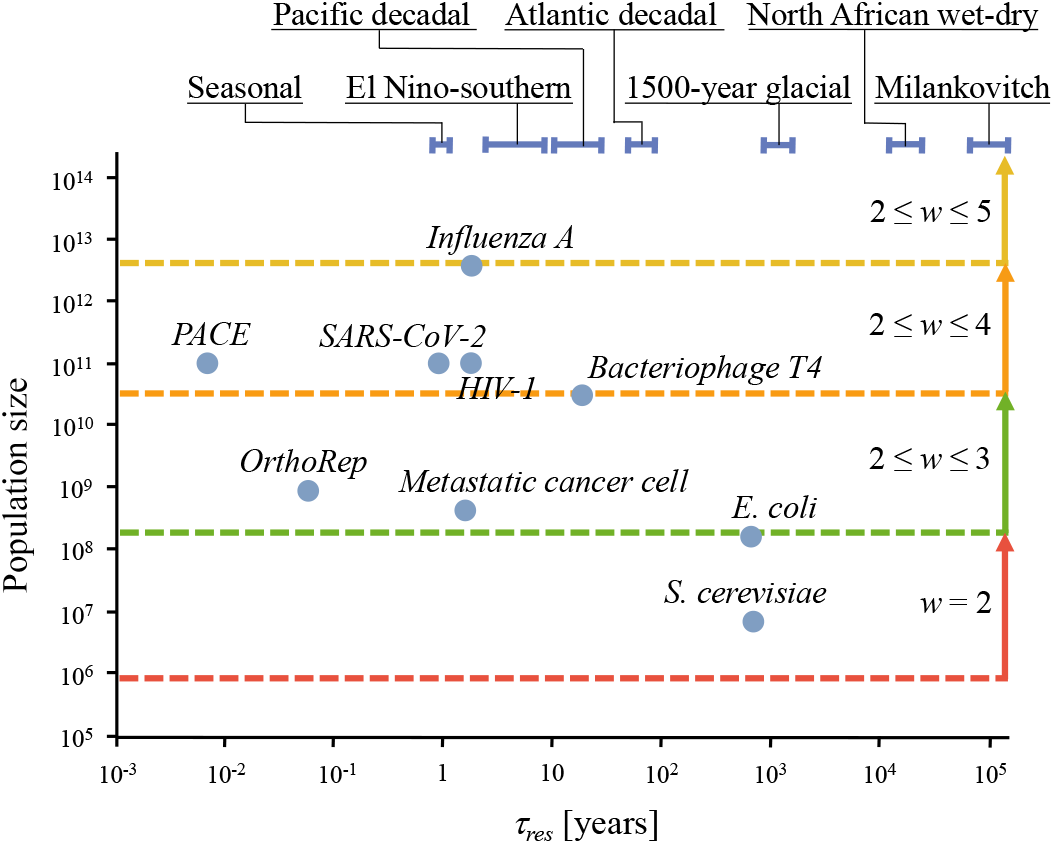
Typical population sizes and resonant times for different organisms. The maximum valley that can be crossed at resonance is independent of the resonance time, but depends on population size. Calculations are done for *L* = 20; different *L* values will inversely rescale the resonance time. Top: recurring environmental fluctuations illustrate processes that could drive a wide range of resonance frequencies.

Figure 3 indicates that *w* = 2 and *w* = 3 valley-crossing can occur on evolutionary timescales if the selection strength fluctuates at resonance for populations larger than 10^6^ − 10^8^. It also demonstrates that traversing a three-step valley (*w* = 4) at resonance would require populations in excess of 10^10^. These data indicate that, assuming bacterial mutation rates have not been much higher than current rates for most of evolutionary history, protein innovation proceeded exclusively by crossing fitness valleys of size 1 and 2.

At short times (*μnt* ≪ 1), we can approximate Eq. 7 by:

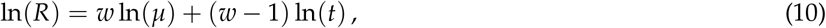

These lines are independent of *n* and *L* (See Fig. 3). The slopes of the lines are equal to the valley size (*w* − 1), and the *y*-intercept is given by *w* ln(*μ*).

#### reversion-controlled regime

If *α >> μ*(*n* − 1)*L*, the resonance time is controlled by the reversion rate back to wild type, and is given by (See Supp. Materials for details):

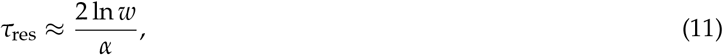

The maximum enhancement of the rate of crossing a fitness valley of length *w* − 1 is given by:

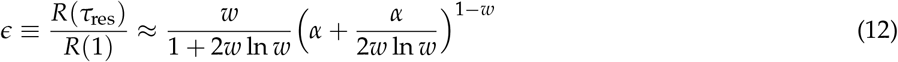

Note that, just as in the diffusion-dominated regime, the rate enhancement grows exponentially with the length of the epistatic valley, with the base of the exponent being controlled by the reversion rate rather than the mutation rate.

### The frequency spectrum of climate oscillations can drive evolutionary innovation

If periods of relaxed selection occur with duration *τ*_res_, the rate of valley crossing is maximal and dependent only on *τ*_res_ and population size. For the mutation-controlled regime, Fig. 4 shows the sizes of valley that can be crossed during one resonant period as a function of *τ*_res_ and population size. A range of genomic hosts that could support protein valley-crossing, including both natural hosts and bioengineering platforms designed for accelerated continuous culture directed evolution (data and references in Suppl. Materials). These data predict that protein valley-crossing in viral and cancer hosts can occur on physiologically relevant timescales. For the majority of proteins that evolved in single-celled organisms, valley-crossing is maximized when fluctuations occur on the millenial time-scale. Although the causes of changing fitness can be system-specific, there exist reliable global oscillations that could drive protein evolutionary resonance across this wide range of timescales (Fig. 4).

## III. Discussion

The non-monotonicity of the valley-crossing rate with diffusion time is reminiscent of recent computational results for alternating selection between two different fitness landscapes in a phenomenological model of immune system adaptation, which found that the fraction of generalists is maximized at intermediate values of alternation frequency [26]. However, in that study, the optimal time between alternations is constrained by the time it takes for the population to equilibrate to a new landscape via selection, and is fundamentally different from the resonance time reported here, which is constrained by the time for genetic information erasure via mutational diffusion.

This work identifies the relevant mutation rates or, equivalently, evolutionary timescales for fluctuating selection to drive valley-crossing. The inability to observe fitness valley crossing via fluctuating selection pressure in directed evolution experiments so far [25, 50] can be explained by the resonance period for proteins in bacteria. From Eq. 8, the resonant frequency for alternating selection in bacteria is three to four orders of magnitude longer than the typical experimental scales. However, the theory predicts that fluctuating selection can significantly enhance valley-crossing for next-generation accelerated directed evolution platforms that bring *τ*_res_ to the timescale of laboratory experiments. In the natural world, fluctuating selection pressures can exponentially accelerate protein optimization on seasonal and evolutionary timescales for viruses and microorganisms, respectively (Fig. 4). Therefore, repeated climate fluctuations over many orders of magnitude in temporal frequency indicates the possibility of repeated bursts of protein optimization that occurred synchronously across a diverse set of hosts. If so, this would represent a essential role of climate fluctuations in driving molecular evolution.

## IV. Supplementary information

### I. Derivation of the valley crossing equation

The set of *L* sites independently experience mutation and weak selection during the mutational diffusion process. The target is to achieve specific point mutations on a subset of *w* sites while keeping the other *L* − *w* sites unchanged. Consider a single site which is capable of being in *n* different states; if the sites are amino acids, *n* = 20. Letting **m**(*t*) be an *n*-dimensional vector such that *m*_*k*_(*t*) is the probability that the site will be in the *k*^th^ state at time *t*. Then, *m*_*k*_(0) = *δ*_*ik*_ and ∑_*k*_ *m*_*k*_(*t*) = 1, where *i* is the state of the site at time 0 and *δ*_*ik*_ is the Kronecker delta function. Without loss of generality, set *i* = 1 to be the identity of the original wild-type state. The probability distribution for the site after time *t* is **m**(*t*) = Θ(*t*) × **m**(0), where the time evolution matrix Θ(*t*) obeys the differential equation:

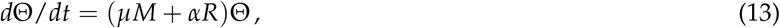

where *μ* is the mutation rate of the organism. At each site, the mutation transfer matrix *M* connects the prior amino acid with the probability of mutating to a new amino acid per generation. For simplicity, only point mutations are considered and the mutation rate amongst all amino acids is assumed to be equal, although *M* can be made to reflect the empirical mutation rates between amino acids as we do in the Results section. *M* is an *n*-by-*n* square matrix:

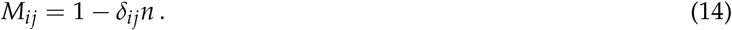

All timescales are in units of number of generations, and multiple simultaneous mutations within a single generation corresponds to the case for which *w >* 1 and *t <*= 1. The reversion, or atavism, transfer matrix *R* accounts for the net transition from all mutants back to the wild type amino acid (denoted without loss of generality by *k* = 1) due to a small selection pressure in favor of the original mutant. Alternatively, in the context of a disconnected population undergoing relaxation of a selection pressure faced by the original population, *R* can represent the extent of coupling between the two populations resulting in constant injection of the original mutant (*k* = 1). In either case, the effective rate of probability flow from a mutation back to the wild type is parameterized by *α*. Thus, the reversion matrix *R* is the *n*-by-*n* square matrix:

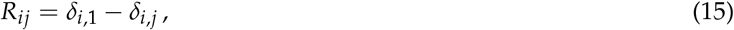

The solution to Eq. 13 is: Θ(*t*) = e^(*μM*+*αR*)*t*^, which is a series expansion of the exponential in powers of the matrix (*μM* + *αR*)*t*. To solve this expansion, we recognize that *M* and *R* constitute, up to scalar multiplication, the right zero semigroup of order two with the following properties: *M*.*M* = − *nM, R*.*M* = − *M, M*.*R* = − *nR*, and *R*.*R* = − *R*:

First, we will prove that the mutation matrix M and the reversion matrix R form a right semigroup. The self-products of the mutation and reversion matrices are:

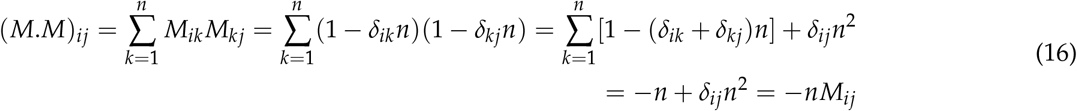

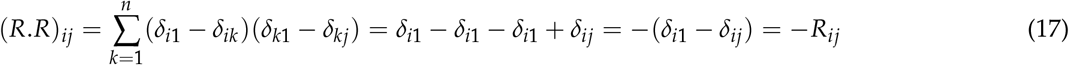

Their mixed products are:

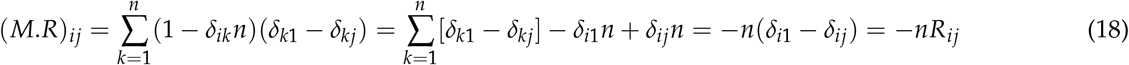

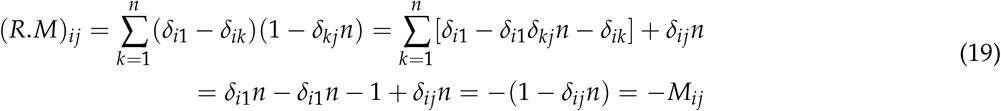

repeatedly applying these identities to linearize the series expansion of Θ(*t*) = e^(*μM*+*αR*)*t*^ and collecting terms yields the time evolution operator as a linear function of *M* and *R*:

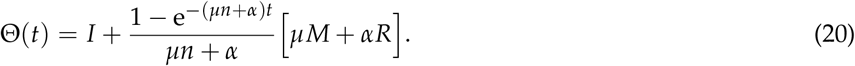

Here, *I* is the identity matrix. Eq. 20 has the expected limits: Θ(0) = *I* and Θ(∞)_*ij*_ = 1/*n* if *α* = 0, corresponding to the identity matrix and the matrix that transforms all initial conditions into the equal likelihood of ending in any mutation, respectively. The insight from Eq. 20 is that multiple consecutive generations of mutation and selection (i.e. reversion to the wildtype) is mathematically equivalent to a single generation in which the mutation and selection matrices are renormalized by the factor 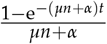. All entries in the first column of Θ(*t*), except the entry on the first row corresponding to the wild-type mutant, are equal to each other. Their value is *p*(*t*) given by Eq. 1 in the main text, which is the probability of achieving any one of the non-wild-type mutants at time *t*.

### II. Universal mutational response curve for *w* = 1

To map all data to a universal function, consider the case for which *α > nμ*. In this case, we have:

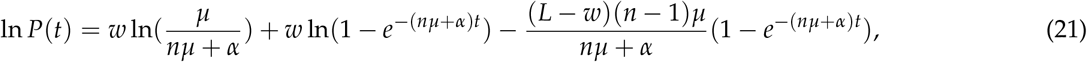

where the last term follows from taking the first term in the series expansion of ln(1 − *x*) for *x <* 1. Then, Eq.6 in the main text follows from the definition of the transformed time and probability variables *t*^*^ and *P*^*^(*t*^*^).

### III. Characteristic time and long time behavior of *P*_*w*_(*t*): *t*_char_ and *N*_latent_

*t*_char_ corresponds to the time of the maximum of *P*_*w*_(*t*). Differentiating the expression for *P*_*w*_(*t*) (Eq. 2 in main text) with respect to *t*, and collecting the only product term that can change signs (note that the *p*(*t*) and 1 − *p*(*t*) terms are non-negative as probabilities, the condition is:

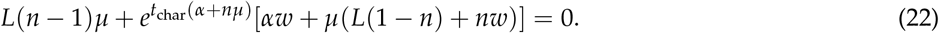

In the diffusion controlled regime, *α << nμ*, so defining a small variable *δ* ≡ *nμ*/*α*, we can expand the solution in powers of *δ*:

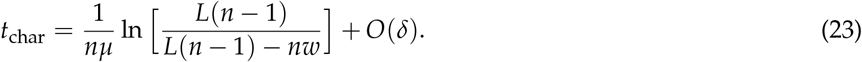

We can further simplify by noting that *L* is typically much larger than *w* to approximate the logarithm as approximately 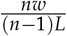 to yield Eq.4 in the main text:

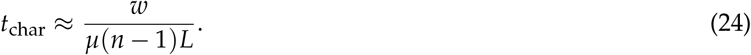

To assess the persister population, consider *P*_*w*_(*t*) the long-time limit (*t* → ∞). To simplify the analysis, define a new variable corresponding to the ratio of reversion to diffusion (i.e. mutation rates): 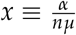. Then:

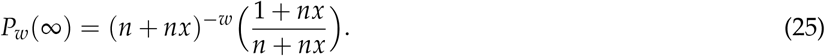

Differentiating with respect to *x* yields the condition for the maximum steady-state persister population:

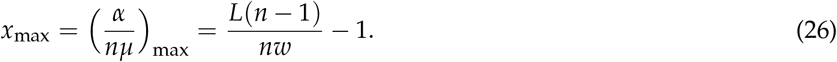

At this optimal condition the persister probability, and hence the maximum persistor probability, is:

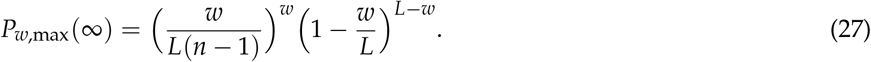

To further simplify, we note that *w* is typically much smaller than *L*, such that 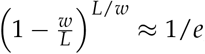, and therefore:

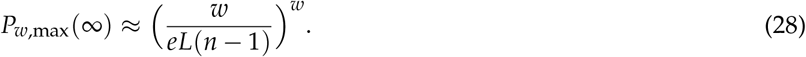

Therefore, the minimum population size to support a persister population, *N*_latent_, is given by the condition *N*_latent_*P*_*w*,max_(∞) = 1, which yields Eq. in the main text:

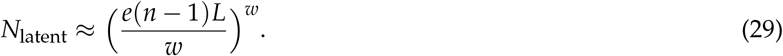

### IV. Resonance time *τ*_res_

*τ*_res_ corresponds to the time of maximum *R*_*w*_(*t*). Differentiating *R*_*w*_(*t*) with respect to *t* and collecting terms, the only factor that can change signs is:

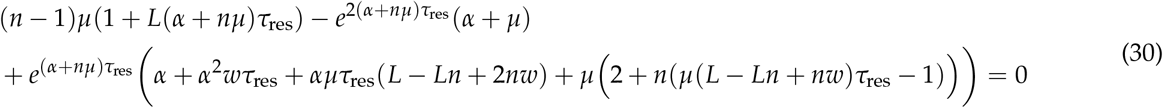

#### IV.1 diffusion controlled regime

In the diffusion controlled regime *α << nμL*, the above simplifies to leading order in *α*/(*nμL*):

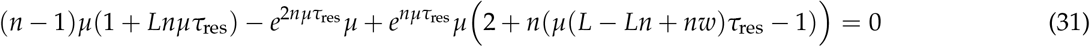

From Eq.4 in the main text, *t*_char_(*n* − 1)*μ* ≈ *w*/*L <<* 1, we can make the ansatz that the same applies to *τ*_res_: *τ*_res_*nμ <<* 1. We will check the resultant prediction of *τ*_res_ to make sure this ansatz is self-consistent. Then, we can expand in powers of *ϵ* ≡ *τ*_res_*nμ* to obtain an expression from the first two orders:

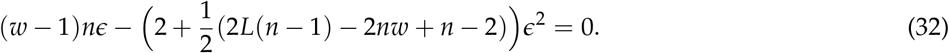

Finally, simplifying using the fact that *w << L*, we obtain the condition:

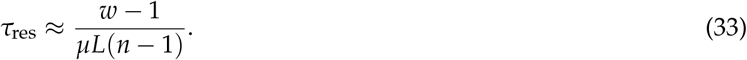

which is consistent with the ansatz that *τ*_res_*nμ <<* 1, and which is Eq. 8 in the main text.

#### IV.2 reversion-controlled regime

In this case, we take the other limit: *λ* ≡ *μn*/*α <<* 1, which yields, to first order in *λ*, the maximal condition:

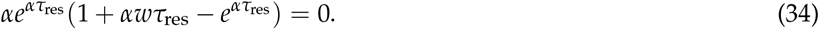

The real solution is:

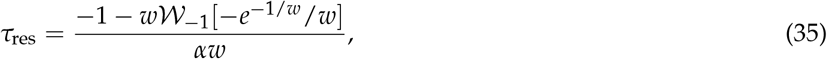

where the function 𝒲_−1_[…] is the -1 branch of the Lambert W function. For simplicity, note that this expression is well approximated by Eq. 11 in the main text:

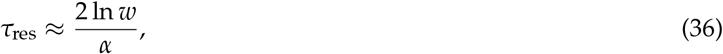

with the approximation error approaching zero as *w* approaches 1.

### V. Simulation methodology for heterogeneous mutation rates

In nature, not all the transitions between amino acids are equally probable, and in fact some transitions are forbidden. As it was stated before, the matrix *M* in Eq. 14 considers that mutation rates among all amino acids are equal. In order to reflect the non-homogeneity of the mutation rates between different amino acids, we need to replace the former matrix *M* with a matrix *P*, where the element *P*_*ij*_ gives the probability that amino acid *i* mutates to amino acid *j*, and *P*_*ii*_ gives the probability that the amino acid *i* does not mutate.

We start this analysis by considering the accepted point mutation matrix (PAM) introduced by Dayhoff et al., which specifies substitutability of amino acids [51]. The base unit of time (uot) of the PAM matrix (a PAM uot) is the time required to change, on the average, 1% of the amino acids, whereas matrix *M* is defined per generation. A PAM uot will depend on many factors, in particular the replication time and the base pair mutation rates of an organism.

To illustrate the effect of the non-homogeneous mutation rates among amino acids, we consider the particular case of the bacterium *E. coli*, whose genome encodes approximately 1.37 × 10^6^ amino acids. The mutation rate of this microorganism was first estimated by Drake *et al*., obtaining 2.5 × 10^−3^ mutations per genome per generation [52]. However, a most recent experiment conducted by Lee *et al*. revealed a value of 10^−3^ mutations per genome per generation [53]. In the same experiment, the aforementioned authors also found that only 69% of the mutations resulted in a modification of the encoding amino acid, being the rest synonymous mutations. Thus, for the *E. coli*, to change 1% of amino acids (1 PAM uot) will require approximately 2 × 10^7^ generations.

We define our matrix *P* = *PAM*^*^ − *I*, where *PAM*^*^ is the PAM matrix expressed in generations and *I* is the identity. Note that, since every element of the matrix *PAM*^*^ represents a probability, the sum over a column must be equal to 1. We can estimate a realistic value of the parameter *μ* in the homogeneous case by solving the equation tr (*P*) = tr (*μM*), which gives 2.58 × 10^−11^ per site per individual per generation.

Numerical simulations of Eqs. 13 and 21 replacing *μM* with matrix *P* and setting *α* = 0 are shown in Fig. 3. The figure shows hundreds of curves simultaneously plotted which correspond to some random combinations of amino acids, for the case *l* = 20. As indicated, different colors represent the different epistatic orders. Dark curves correspond to the analytical result of the homogeneous case given by Eq. 2 and the above-said value of the mutation rate. We can see an excellent agreement between the homogeneous and the heterogeneous cases, laying the homogeneous curves in the middle of the spectrum, which is approximately 2 orders of magnitude thick.

### VI. Fitting the theory to data in Figure 2

Data sets were taken from Ref. [42–45] and were modified from the original figures by normalizing the probability to its maximum. The data sets were fitted by Eq. 2 (solid lines), setting *w* = 1, *n* = 4 and *α, L* and *μ* as free parameters. The value of the free parameters obtained are: (A) *α* = 1.49 × 10^−5^, *μ* = 1.87 × 10^−3^ and *L* = 118; (B) *α* = 3.51 × 10^−6^, *μ* = 7.86 × 10^−4^ and *L* = 112; (C) *α* = 3.30 × 10^−7^, *μ* = 3.87 × 10^−4^ and *L* = 136; (D) *μ* = 3.86 × 10^−3^, *α* = 0.13 and *L* = 135; (E) *μ* = 2.54 × 10^−3^, *α* = 0.054 and *L* = 130; (F) *μ* = 2.25 × 10^−3^, *α* = 0.153 and *L* = 120.

#### Eq. 5 in Ref [42] as a special case of Eq. 2

It is evident from the previous fittings that figures A-C correspond to the case where *α* = 0. Since *n* ∝ *t* in Eq. 5 in Ref [42], we can identify this equation as a special case of our Eq. 2, setting *w* = 1 and *α* = 0. In the limit where *μnt* ≪ 1, the correspondence between the parameters of both equations is given by *α*^′^ = (*L* − 1)(*n* − 1)*μ* and *f*_+_ = *μ*.

Dashed lines correspond to Eq. 5 in Ref [42] with *α*^′^ and *f*_+_ given by the above-mentioned relationships and the values of the parameters obtained by the fitting curves in each case.

For panel (G), all data sets were mapped using the transformations 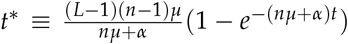 and *P*^*^ ≡ (*n* − 1)(*L* − 1)*P*, with the parameter values from each of the fitting curves and then plotted following Eq. 5.

### VII. Estimating resonance times for different hosts

The data for effective population size and mutation rates for organisms and directed evolution experimental platforms shown in Fig.4 are summarized in Table S1.

**Table S1:** Data for resonance times calculated in Figure 4. * Number of CD4+ T cells infected times viral burst size.** Effective population size of E. coli times viral burst size

| Organism | $\mu$(per bp per gen) | Generation time | Population size [54] | Reference |
| --- | --- | --- | --- | --- |
| OrthoRep | $10^{-5}$ | 120 min | $10^9$ | [33] |
| PACE | $5 \times 10^{-5}$ | 48 min | $10^{11}$ | [55] |
| HIV-1 | $3.4 \times 10^{-5}$ | 52 hr | $10^{11} *$ | [56] |
| Influenza A | $2 \times 10^{-6}$ | 7 hr | $10^{12} *$ | [57] |
| Sars-CoV-2 | $5.7 \times 10^{-6}$ | 24 hr | $10^{11} *$ | [58] |
| Bacteriophage T4 | $2.4 \times 10^{-8}$ | 60 min | $3.6 \times 10^{10} **$ | [52] |
| Escherichia Coli | $2.2 \times 10^{-10}$ | 30 min | $1.8 \times 10^8$ | [53] |
| Saccharomyces cerevisiae | $3.8 \times 10^{-10}$ | 90 min | $8 \times 10^6$ | [59] |
| Metastatic cancer cell | $5 \times 10^{-6}$ | 28 hr | $10^8 - 10^9$ | [60] |

## Notes

### Competing Interest Statement

The authors have declared no competing interest.

